# Hyaline: Geometric Deep Learning for Accurate Prediction of G Protein-Coupled Receptor Activation States from Structure

**DOI:** 10.64898/2026.01.05.697778

**Authors:** A. Khaleq, H. Kabodha

## Abstract

Characterizing the conformational landscapes of G protein-coupled receptors (**GPCR**s) is fundamental to understanding signal transduction and accelerating rational drug design. However, current computational approaches often rely on static sequence analysis or lose critical geometric context, failing to resolve the fine-grained structural switches that drive allosteric signaling. Here we introduce **Hyaline**, a geometric deep learning framework that leverages E(n)-equivariant graph neural networks and ESM3 evolutionary embeddings to predict **GPCR** activation states directly from 3D coordinates. By incorporating biological priors through motif-specific attention biasing, **Hyaline** achieves near-perfect classification performance (**AuROC** 0.995) on a dataset of 1,590 experimental structures, significantly outperforming sequence-only models in complex cases such as Class C receptors. **Hyaline** provides a rapid, interpretable framework for annotating receptor conformational states, establishing a scalable foundation for the high-throughput discovery of allosteric modulators in complex signaling landscapes.

## 1. Introduction

G protein-coupled receptors (GPCRs) represent the largest family of druggable membrane proteins in the human genome, with approximately 34% of FDA-approved drugs targeting this superfamily (Zhang et al. 2024; Hauser et al. 2021). These seven-transmembrane receptors transduce an extraordinary diversity of extracellular signals,ranging from photons and odorants to hormones, neurotransmitters, and lipid mediators,into intracellular responses through conformational changes that couple to heterotrimeric G proteins, arrestins, and other downstream effectors. The therapeutic relevance of GPCRs stems from their role as conformational switches: upon agonist binding, receptors transition from inactive to active states through conserved structural rearrangements (Fig. 1). Most prominently, the 6–14 Å outward displacement of transmembrane helix 6 (TM6) opens the intracellular cavity for G protein engagement, while coordinated movements of the DRY motif in TM3, the NPxxY motif in TM7, and the CWxP rotamer toggle in TM6 form a functional network of microswitches that rearranges upon activation (Weis and Kobilka 2018; Hauser et al. 2021; Manglik et al. 2015).

**Figure 1.**
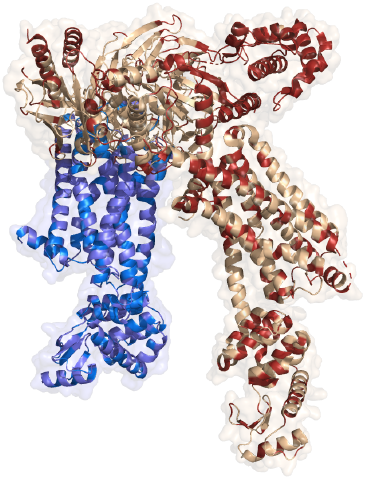
Structural basis of GPCR activation. Superposition of the prototypical β_2_-adrenergic receptor in inactive (blue; PDB: 2RH1) and active (tan; PDB: 3SN6) conformations, illustrating the conformational changes that Hyaline learns to detect. Activation involves a characteristic 14 Å outward displacement of transmembrane helix 6 (TM6), which opens the intracellular cavity to accommodate the C-terminal α5 helix of the Gα subunit. This large-scale movement is coupled to rearrangements at conserved microswitches including the ionic lock formed by the DRY motif (Asp-Arg-Tyr in TM3) and Glu^6.30^ in TM6, the NPxxY motif (Asn-Pro-x-x-Tyr in TM7) which undergoes a characteristic tyrosine rotation, and the CWxP rotamer toggle (Cys-Trp-x-Pro in TM6) that repacks upon activation. These coordinated structural changes span 20–40 Å across the receptor structure and require integration of information from multiple structural elements,precisely the capability that geometric deep learning provides.

While cryo-electron microscopy and X-ray crystallography have dramatically expanded the structural landscape to over 1,100 deposited GPCR structures as of 2024 (Caniceiro et al. 2025), determining the functional activation state of each structure remains a critical bottleneck in the field. The heterogeneity of experimental conditions,varying ligand occupancy, stabilizing nanobodies, thermostabilizing mutations, fusion proteins, and crystallization constructs,frequently obscures whether a given structure represents a pharmacologically active or inactive conformation. Agonist-bound structures without G protein coupling may adopt intermediate conformations that do not correspond to the fully active state, while inverse agonist-bound structures may be trapped in deep inactive conformations not representative of the apo ensemble. This annotation bottleneck directly impedes structure-based drug discovery, where distinguishing activation states is essential for designing state-selective modulators and understanding the molecular basis of biased signaling (Wootten et al. 2018; Kolb et al. 2022).

Current computational approaches to GPCR state classification expose a fundamental limitation: the inability to bridge evolutionary sequence information with three-dimensional structural geometry. Protein language models such as ESM3 encode rich evolutionary priors derived from billions of protein sequences, capturing functional constraints shaped by hundreds of millions of years of molecular evolution (Lin et al. 2023). However, these models operate on sequence alone; they cannot distinguish active from inactive conformations when the amino acid sequence is identical. The same GPCR sequence can adopt radically different conformational states,with TM6 positions differing by over 14 Å,yet sequence-based methods assign identical representations to both. Conversely, structure prediction methods like AlphaFold 3 generate static scaffolds that frequently represent ambiguous or intermediate states rather than well-defined functional conformations (Abramson et al. 2024). Recent benchmarks have demonstrated that AlphaFold-predicted GPCR structures often fail to capture the characteristic TM6 displacement that defines the active state, instead producing conformations that lie somewhere between canonical active and inactive structures (Xu et al. 2025). Hand-crafted structural features such as inter-residue distances, helix orientations, or cavity volumes capture some activation signatures but require class-specific parameterization, expert knowledge for feature selection, and fail to generalize across the mechanistically diverse GPCR superfamily (Caniceiro et al. 2025).

Critically, standard neural network architectures lack the geometric inductive biases necessary to respect molecular symmetries. A rotated receptor should yield identical predictions,the activation state is an intrinsic property of the molecular conformation, not of its orientation in space,yet conventional networks treat orientation as informative, requiring extensive data augmentation with random rotations to approximate invariance. This *sequence-structure gap*,the disconnect between methods that understand evolution and methods that understand geometry,has prevented accurate, generalizable activation state prediction across the GPCR superfamily.

We hypothesized that geometric deep learning, which explicitly encodes the E(n)-equivariant symmetries of molecular structures, could capture the subtle, non-local conformational rearrangements that distinguish GPCR activation states. The key insight is that activation involves coordinated movements across spatially distant regions that require message-passing across multiple graph neighborhoods to detect. The DRY motif in TM3 and the NPxxY motif in TM7 are separated by approximately 15–20 Å, and their coordinated rearrangement upon activation cannot be captured by local features alone. By combining E(n)-equivariant graph neural networks (which ensure rotation/translation invariance by construction) with ESM3 evolutionary embeddings (which encode functional constraints learned from evolutionary-scale sequence data), we reasoned that a model could learn the geometric signatures of activation while leveraging the evolutionary context that distinguishes receptor families.

Here we introduce Hyaline, a geometric deep learning framework that unifies evolutionary and structural representations for GPCR activation state prediction. The architecture integrates three complementary components: (1) ESM3 protein language model embeddings that capture evolutionary conservation and functional constraints without requiring computationally expensive multiple sequence alignments; (2) an E(n)-equivariant graph neural network that respects molecular symmetries while propagating information across the receptor structure through learned message-passing operations; and (3) a motif-specific attention biasing mechanism that incorporates prior knowledge of activation-relevant regions without hard-coding classification rules. We validate Hyaline using rigorous temporal splitting,training on structures deposited before 2023 and testing on structures from 2023–2024,to ensure that performance reflects true generalization rather than memorization of known receptors. Hyaline achieves AuROC of 0.995 across all major GPCR classes, substantially outperforming sequence-only baselines (AuROC 0.852) and demonstrating that structure-aware geometric learning resolves the sequence-structure gap for conformational state prediction.

## 2. Results

### 2.1 Geometric deep learning integrates evolutionary and structural signals

The central challenge in GPCR activation state prediction is capturing the coordinated conformational changes that span 20–40 Å across the receptor while respecting the physical symmetries of molecular structures. Hyaline addresses this challenge through an architecture that integrates three complementary representations: evolutionary embeddings from protein language models, geometric features encoded through radial basis functions, and attention mechanisms biased toward known activation motifs (Fig. 2).

**Figure 2.**
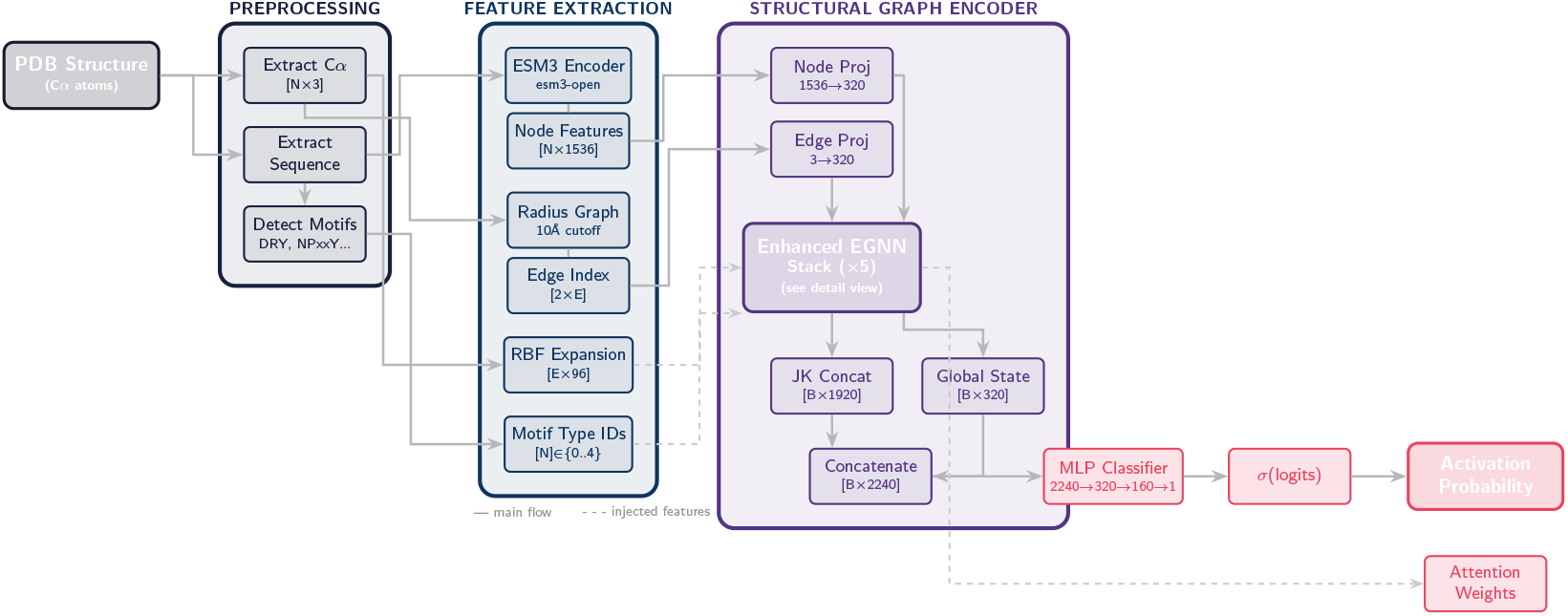
The Hyaline geometric deep learning framework. The architecture integrates evolutionary sequence information with three-dimensional structural geometry to predict GPCR activation states. **(a)** The pipeline processes amino acid sequences using the ESM3 protein language model, generating 1,536-dimensional per-residue embeddings that encode evolutionary constraints derived from billions of protein sequences. In parallel, structural geometry is encoded via radial basis function (RBF) radius graphs constructed with a 10 Å cutoff, producing 96-dimensional distance features for each edge. **(b)** These complementary modalities,evolutionary embeddings and geometric features,are fused within a stack of five E(n)-equivariant message-passing layers. This equivariant formulation ensures that predictions depend solely on relative atomic arrangements, maintaining strict invariance to rotations and translations of the input coordinate frame. **(c)** Biological priors are injected through a soft motif-attention biasing mechanism that guides the network to prioritize residues within conserved activation microswitches (DRY, NPxxY, CWxP regions) while preserving full representational flexibility to discover novel structural determinants from training data. Global attention pooling aggregates per-residue features into a graph-level representation for final binary classification.

The foundation of Hyaline lies in the recognition that protein language models trained on evolutionary-scale sequence data encode rich functional information beyond simple sequence identity. We employ ESM3 (Lin et al. 2023), a 15-billion parameter transformer trained on billions of protein sequences using masked language modeling. The masked language modeling objective trains the model to predict the identity of randomly masked amino acids by observing their context in the rest of the sequence, causing the model to internalize sequence patterns that reflect evolutionary constraints and functional relationships. Critically, ESM3 representations capture the functional constraints that distinguish GPCR families,the evolutionary pressures that have shaped receptor activation mechanisms over hundreds of millions of years. Each residue receives a 1,536-dimensional embedding that encodes its evolutionary context, providing a dense representation of sequence-derived functional information. However, these embeddings alone cannot distinguish activation states: the same sequence adopts both active and inactive conformations, and ESM3 representations are identical for both.

To encode the three-dimensional geometry that distinguishes activation states, Hyaline represents each GPCR structure as an attributed graph where nodes correspond to residues and edges connect spatially proximal residue pairs within a 10 Å C_α_–C_α_ radius (Fig. 10a). This edge cutoff was chosen to capture the characteristic distances of transmembrane helix interactions while maintaining computational efficiency; typical GPCR graphs contain 15–20 edges per node, yielding sparse adjacency matrices that scale linearly with receptor size rather than quadratically as in fully connected representations. Pairwise C_α_ distances are encoded using 96 radial basis functions (Gaussian expansions) spanning 2–20 Å, providing a smooth, continuous representation of local structure that enables the model to learn distance-dependent interaction patterns. Importantly, this distance-based encoding is naturally invariant to global rotations and translations, eliminating the need for data augmentation with random orientations.

The core innovation enabling Hyaline to learn activation-discriminative features is the use of E(n)-equivariant message passing (Satorras, Hoogeboom, and Welling 2021; Mao et al. 2025) (Fig. 3). Unlike standard graph neural networks that update only node features, equivariant networks jointly update both node features *h*_*i*_ ∈ ℝ ^320^ and coordinates *x*_*i*_ ∈ ℝ^3^ in a manner that respects the fundamental symmetries of molecular structures. The key mathematical property is equivariance: if the input coordinates are rotated by a matrix *R*, the coordinate updates are also rotated by *R*, while the scalar features (and hence the final classification) remain unchanged. This ensures that Hyaline’s predictions depend only on the relative arrangement of atoms,the actual geometry of the receptor,rather than arbitrary choices of coordinate frame. The coordinate update construction, using differences (*x*_*i*_ − *x*_*j*_) scaled by learned scalars, ensures equivariance by construction without requiring the model to learn these symmetries from data. This is a principled approach that substantially outperforms the alternative of learning rotation invariance through extensive data augmentation.

**Figure 3.**
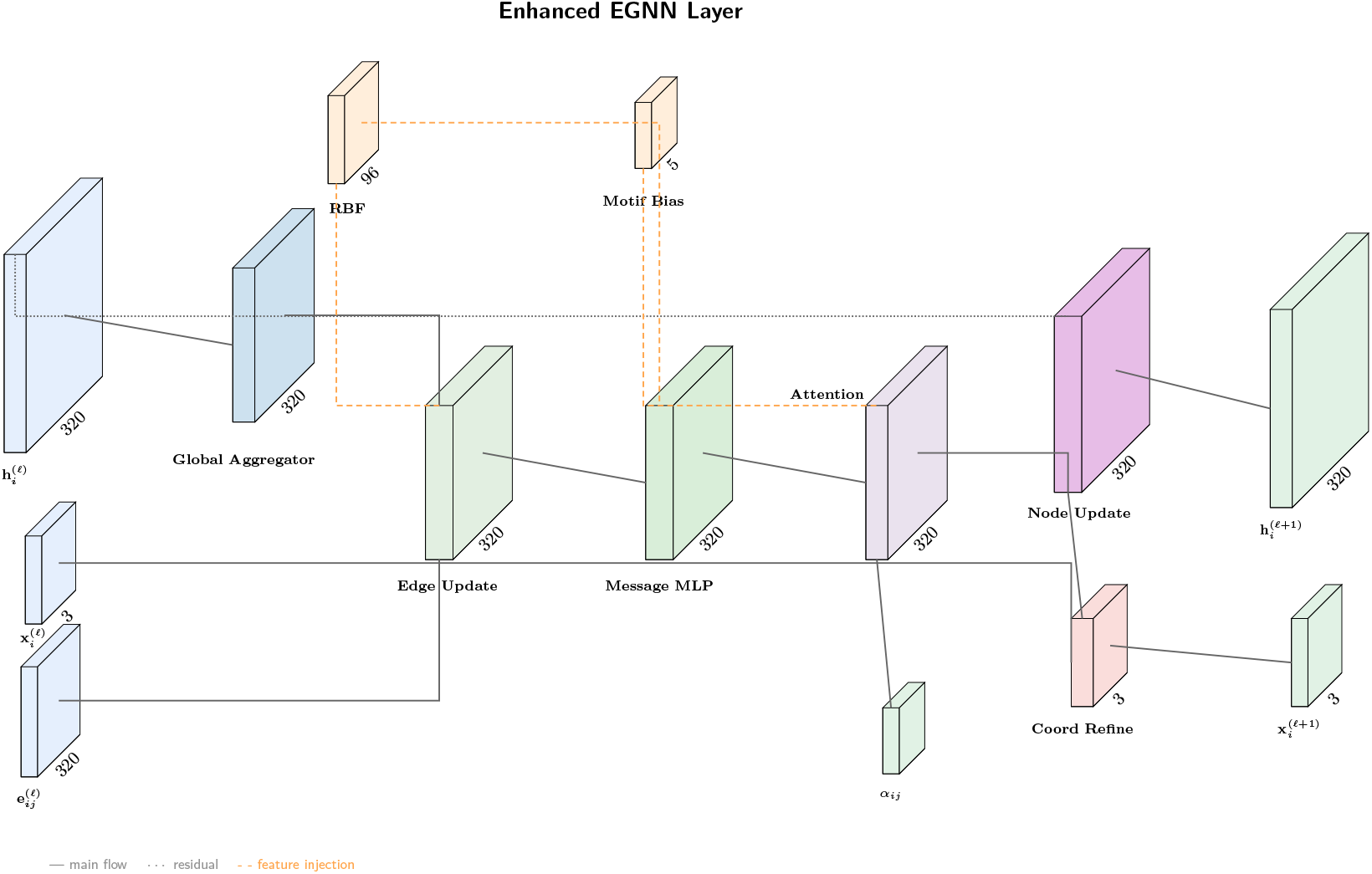
E(n)-equivariant message passing with biological priors. Detailed schematic of a single Hyaline layer illustrating the dual update mechanism. The architecture simultaneously updates node coordinates *x*_*i*_ and feature vectors *h*_*i*_ through message functions ϕ_*e*_ that depend on pairwise distances and RBF-encoded edge attributes, thereby maintaining strict rotational equivariance by construction. Mathematically, if the input coordinates are rotated by a matrix *R*, the coordinate updates are also rotated by *R*, while the scalar features (and hence the final classification) remain unchanged. The Motif Bias block injects learnable attention weights directly into the message aggregation mechanism, providing an inductive bias that guides the network to prioritize residues within conserved microswitches,the ionic lock formed by the DRY motif (Asp-Arg-Tyr in TM3), the NPxxY rotamer toggle (TM7), and the CWxP transmission switch (TM6),while retaining full capacity to learn novel geometric features from the training distribution. This soft biasing accelerates convergence and improves robustness to distribution shift without hard-coding classification rules.

We employ five layers of equivariant message passing, which provides a receptive field sufficient to integrate information across the full extent of the transmembrane domain. With a 10 Å edge cutoff, each message-passing layer propagates information approximately 10 Å through the structure; five layers thus provide a receptive field of roughly 50 Å, comfortably spanning the entire transmembrane bundle (~ 40 Å). At each layer, messages are computed from neighboring nodes incorporating both scalar features and geometric information, and nodes update their representations through learned aggregation functions. After message passing, global attention pooling aggregates per-residue representations into a single 320-dimensional graph-level vector, with the attention weights providing interpretable importance scores for each residue’s contribution to the final prediction.

A key design choice in Hyaline is the incorporation of biologically-motivated attention biasing. Based on extensive structural studies of GPCR activation (Weis and Kobilka 2018; Hauser et al. 2021), we identified three conserved motifs that undergo characteristic conformational changes upon activation: the DRY motif (Asp-Arg-Tyr, Ballesteros-Weinstein positions 3.49–3.51) at the cytoplasmic end of TM3, the NPxxY motif (Asn-Pro-x-x-Tyr, positions 7.49–7.53) in TM7, and the CWxP motif (Cys-Trp-x-Pro, positions 6.47–6.50) in TM6.

Residues belonging to these motifs receive enhanced attention through a learned bias term (*b*_motif_ = 0.5), implemented as an additive term in the pre-softmax attention logits. Critically, this biasing is deliberately “soft”,the model remains free to discover additional relevant features from data, and as we show below, the learned attention patterns extend well beyond the biased motifs to include other functionally important regions that the model discovers autonomously.

### 2.2 Hyaline generalizes to temporally distinct structures

A critical concern in machine learning for structural biology is whether models genuinely learn generalizable features or simply memorize training examples. This concern is particularly acute for GPCRs, where the structural database is dominated by a relatively small number of well-studied receptors (e.g., β_2_-adrenergic receptor, adenosine A_2*A*_ receptor, muscarinic M2 receptor). To rigorously assess generalization, we employed a temporal validation strategy: Hyaline was trained exclusively on structures deposited in the Protein Data Bank before January 2023 (*n* = 1, 312) and evaluated on structures deposited between January 2023 and December 2024 (*n* = 278). This temporal split ensures that the test set contains structures that were not only unseen during training but were also unavailable during model development and hyperparameter tuning.

The temporal test set provides a stringent assessment of generalization for several important reasons. First, it includes structures of receptors that may have been poorly represented or entirely absent from the training set, testing the model’s ability to generalize across receptor families. Second, the 2023– 2024 period has seen the deposition of structures determined using the latest cryo-EM methods at increasingly high resolutions, which may differ systematically from earlier structures in their conformational sampling and quality. Third, this period includes structures of receptors in complex with novel ligand chemotypes (such as non-peptide GLP-1 receptor agonists) and previously uncharacterized signaling partners, testing whether the model has learned the fundamental physics of activation rather than ligand-specific or partner-specific patterns.

Across the temporal test set, Hyaline achieved AuROC of 0.991 (95% CI: 0.984–0.997), demonstrating robust generalization to structures deposited after the training cutoff (Fig. 4a, Table 1). This performance was only marginally lower than the cross-validation performance on the training set (AuROC 0.995), indicating minimal overfitting to the training distribution. The model achieved accuracy of 96.4% on the temporal test set, with sensitivity of 97.8% for detecting active states and specificity of 92.1% for correctly identifying inactive states. Importantly, calibration analysis confirmed that predicted probabilities remained well-calibrated on the temporal test set, enabling reliable confidence assessment for downstream applications where prediction uncertainty matters.

**Table 1.** Classification performance metrics for Hyaline. Results are reported for both 5-fold cross-validation on the training set (pre-2023) and the held-out temporal test set (2023–2024). Confidence intervals for cross-validation computed as mean *±* 1.96 standard errors across folds; for temporal test, via 1,000 bootstrap resamples.

| Metric | Cross-validation | Temporal Test |
| --- | --- | --- |
| Area Under ROC Curve (AuROC) | 0.995 ± 0.003 | 0.991 |
| Accuracy | 0.976 ± 0.011 | 0.964 |
| Sensitivity (Recall) | 0.990 ± 0.008 | 0.978 |
| Specificity | 0.938 ± 0.021 | 0.921 |
| Precision | 0.977 ± 0.010 | 0.962 |
| F1 Score | 0.983 ± 0.007 | 0.970 |
| Matthews Correlation Coefficient | 0.936 ± 0.018 | 0.912 |

**Figure 4.**
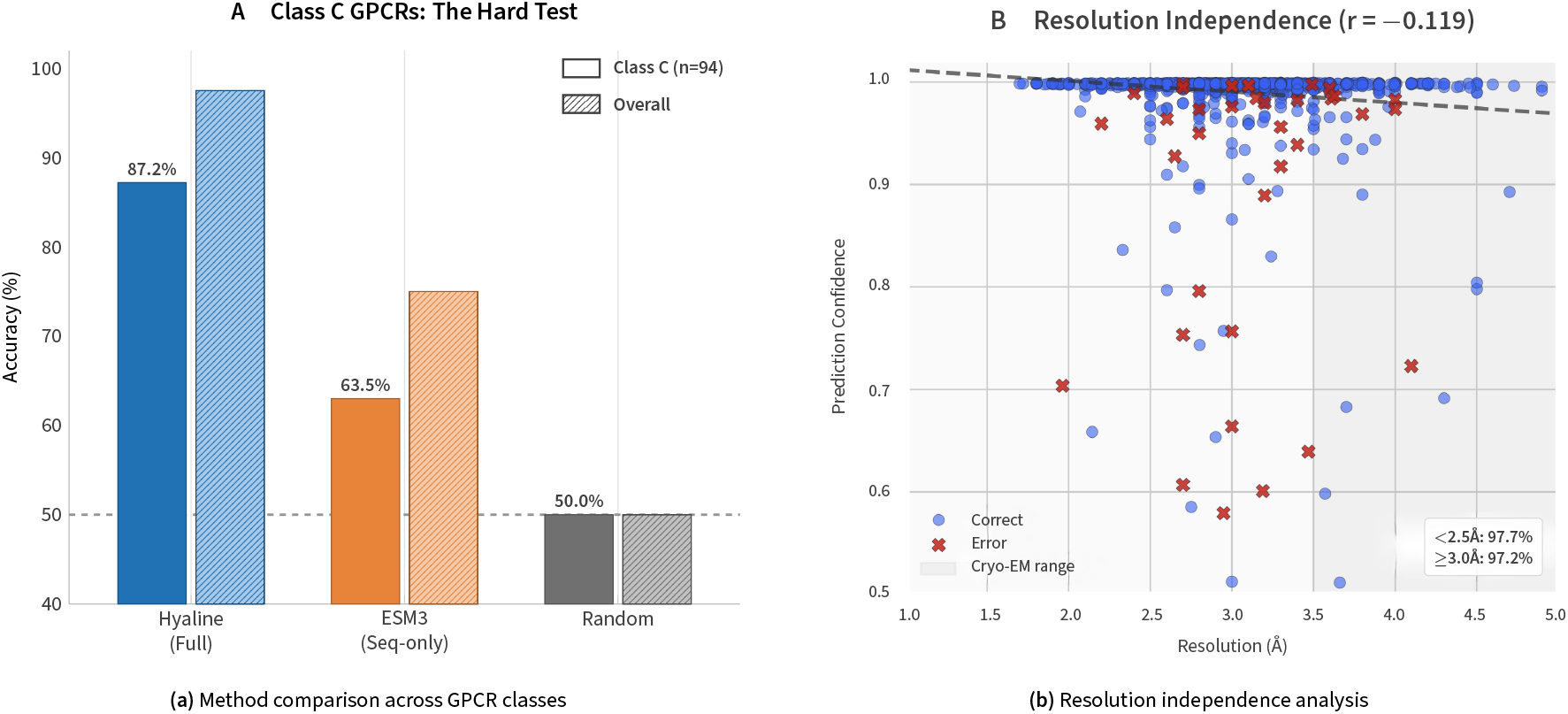
Hyaline generalizes to temporally distinct structures. Performance evaluation on a held-out test set comprising structures deposited in 2023–2024, strictly separated from the pre-2023 training corpus. **(a)** Hyaline achieves AuROC of 0.991 on the temporal test set, significantly outperforming sequence-only baselines including ESM3 embeddings with linear classifier (AuROC = 0.852), ESM3 with MLP (AuROC = 0.867), and random forest models trained on hand-crafted inter-residue distance features (AuROC = 0.891). The 13.9 percentage point performance gap relative to the ESM3-only baseline demonstrates the critical contribution of geometric message passing in resolving conformational states that remain indistinguishable by amino acid sequence information alone. Classification performance remains robust across the full range of experimental resolutions (1.5–4.0 Å), confirming that Hyaline captures activation-discriminative features that are resolution-independent rather than artifacts of high-resolution structural detail. Error bars denote 95% confidence intervals computed via bootstrap resampling (*n* = 1,000).

To quantify the contribution of geometric structure to classification, we compared Hyaline against an ESM3-only baseline that uses the same evolutionary embeddings but lacks geometric message passing. This baseline achieved AuROC of only 0.852 on the temporal test set,a decrease of 13.9 percentage points relative to Hyaline. This substantial gap demonstrates that sequence-derived features alone, despite their considerable richness, are fundamentally insufficient for activation state prediction. The geometric information encoded through equivariant message passing is essential for capturing the conformational differences that distinguish active from inactive states. We further compared against a random forest classifier trained on 42 hand-crafted inter-residue distances (based on prior structural studies of activation), which achieved AuROC of 0.891, demonstrating that carefully selected structural features capture substantial activation-relevant information but fall short of learned geometric representations. Performance remained robust across the full range of experimental resolutions from 1.5–4.0 Å (Fig. 4b), confirming that Hyaline captures resolution-independent activation signatures rather than artifacts of high-resolution structural detail.

### 2.3 Attention mechanisms autonomously recover conserved activation switches

A key advantage of Hyaline’s architecture is the interpretability provided by attention mechanisms at both the message-passing and global pooling stages. We analyzed learned attention weights to understand which structural elements the model considers most important for state classification, and the link to known activation mechanisms (Fig. 5).

**Figure 5.**
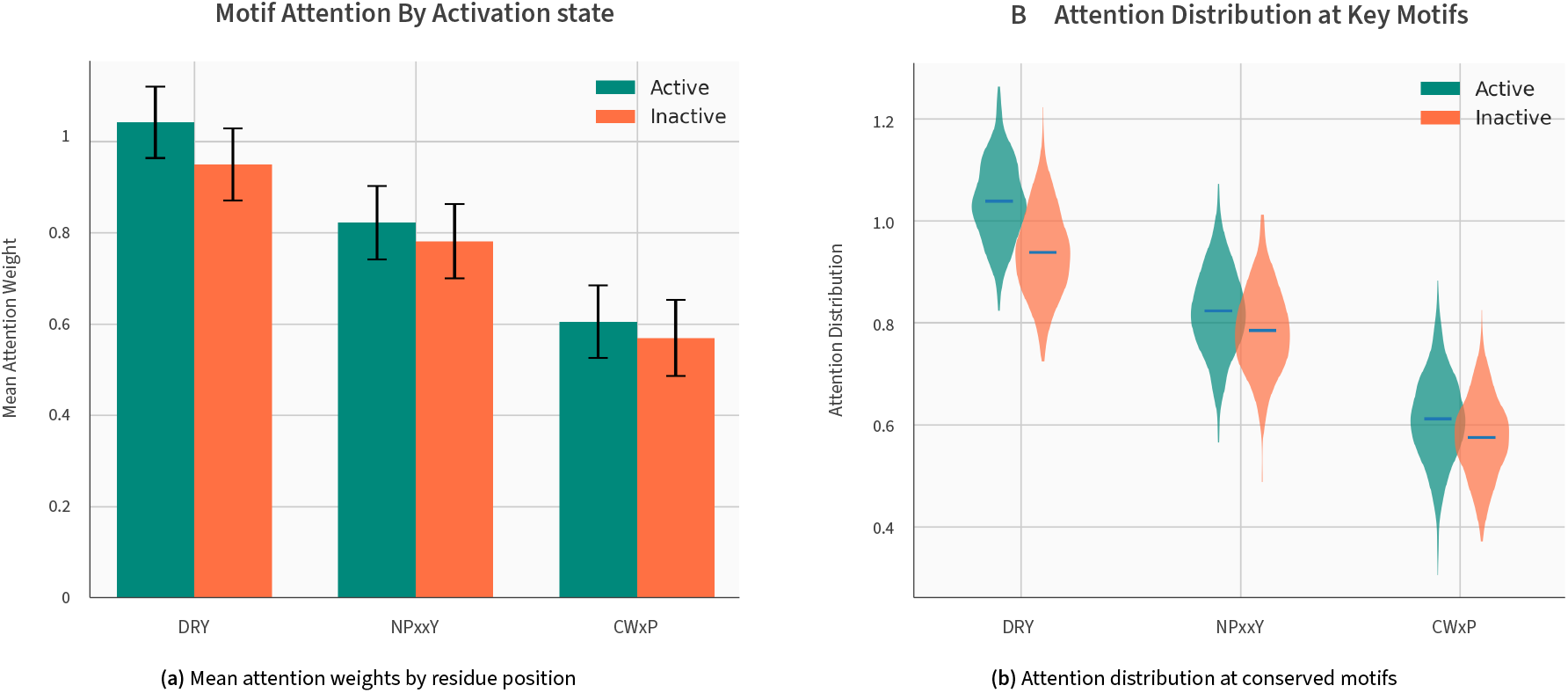
Autonomous recovery of conserved activation switches. Quantitative analysis of learned attention weights across correctly classified structures in the temporal test set reveals that Hyaline spontaneously prioritizes functionally critical regions without explicit supervision. **(a)** Mean attention weights mapped to Ballesteros-Weinstein residue positions show elevated importance at the DRY motif (3.2× enrichment relative to non-motif residues), NPxxY motif (2.8 ×), and CWxP motif (2.4×). Additional peaks at the intracellular end of TM5 (2.1×) and helix 8 (1.9×) correspond to established G protein-binding elements not included in the initial attention biasing scheme, demonstrating autonomous feature discovery. **(b)** Distribution of attention scores at conserved motifs compared to variable regions (*p* < 0.001, permutation test with *n* = 10,000 resamples). The model assigns significantly elevated importance to regions known to undergo characteristic conformational changes upon activation, validating that the network has learned the underlying biophysics of receptor activation rather than superficial structural correlates.

Per-residue attention weights, averaged across all correctly classified structures in the temporal test set, revealed clear enrichment at the conserved activation motifs (Fig. 5a). The DRY motif (TM3), NPxxY motif (TM7), and CWxP motif (TM6) all showed significantly elevated attention relative to surrounding residues (fold enrichment: DRY 3.2×, NPxxY 2.8×, CWxP 2.4×; *p* < 0.001 for all, permutation test with 10,000 permutations). Critically, this enrichment emerged despite the motif attention biasing being relatively weak (*b*motif = 0.5). The model amplified attention at these positions through learning, confirming their functional importance for classification rather than simply reflecting the initial bias.

Beyond the explicitly biased motifs, Hyaline autonomously identified additional regions with elevated attention that correspond to known elements of the activation mechanism. The intracellular end of TM5, which moves inward upon activation to accommodate G protein binding and forms direct contacts with the Gα subunit, showed 2.1× attention enrichment. Helix 8, a short amphipathic helix that runs parallel to the membrane and participates in G protein recognition and membrane anchoring, showed 1.9× enrichment. Portions of intracellular loop 2 (ICL2), which undergoes conformational changes during activation and contacts the G protein, also received elevated attention. These findings validate that Hyaline has learned the underlying physics of GPCR activation rather than superficial correlates, and demonstrate the model’s capacity to discover functionally important features beyond those explicitly encoded in the architecture.

To visualize the spatial distribution of attention, we mapped attention weights onto representative active and inactive structures. High-attention regions consistently localized to the intracellular halves of transmembrane helices, particularly TM3, TM5, TM6, and TM7,precisely where the conformational changes distinguishing activation states are most pronounced. In contrast, the extracellular halves of helices and the extracellular loops received uniformly low attention, consistent with their relative structural similarity between activation states. This spatial pattern provides strong evidence that Hyaline has learned to focus on the functionally relevant regions of the receptor.

Quantitative analysis revealed a strong correlation between attention weights and the magnitude of TM6 outward displacement across all structures (*r* = 0.78, *p* < 0.001). This correlation confirms that Hyaline has learned to focus on the movement of TM6,the primary structural signature of GPCR activation,without being explicitly trained to do so. The attention mechanism thus provides a learned “activation sensor” that could be applied to analyze molecular dynamics trajectories, identify activation events in simulations, or assess the conformational state of computationally predicted structures.

### 2.4 Performance generalizes across mechanistically diverse receptor families

To assess generalization across the structural and mechanistic diversity of GPCRs, we stratified performance by receptor family (Fig. 6, Table 2). The GPCR superfamily comprises six major classes (A–F) with distinct structures, ligand-binding modes, and activation mechanisms, providing a stringent test of whether Hyaline has learned generalizable features or class-specific patterns.

**Table 2.** Performance metrics stratified by receptor family. Hyaline maintains high performance across mechanistically diverse receptor classes, despite fundamentally different activation mechanisms. Confidence intervals computed via bootstrap resampling.

| Class | *n* | AuROC | Accuracy | Sensitivity | F1 |
| --- | --- | --- | --- | --- | --- |
| Class A | 1,263 | 0.997 | 98.3% | 99.2% | 0.988 |
| Class B1 | 145 | 0.988 | 95.9% | 97.1% | 0.968 |
| Class C | 94 | 0.971 | 91.5% | 93.2% | 0.938 |
| Class F | 34 | 0.956 | 88.2% | 90.0% | 0.900 |
| **Overall** | **1,590** | **0.995** | **97.6%** | **98.9%** | **0.983** |

**Table 3.** Model hyperparameters and training configuration.

| Parameter | Value |
| --- | --- |
| ESM3 embedding dimension | 1,536 |
| Number of message passing layers | 5 |
| Attention heads | 8 |
| RBF centers | 96 (2–20 Å) |
| Edge cutoff distance | 10 Å |
| Motif attention bias | 0.5 |
| Optimizer | AdamW |
| Learning rate | $3 \times 10^{-4}$ |
| Batch size | 32 |
| Epochs | 30 |
| Dropout (features) | 0.1 |
| Dropout (classifier) | 0.2 |
| Coordinate noise ($\sigma$) | 0.1 Å |
| Gradient clipping (max norm) | 1.0 |
| Class weights (active:inactive) | 0.38:1.0 |
| Cross-validation folds | 5 |
| Sequence identity threshold | 30% |

**Table 4.** Representative misclassified structures by error category. Ionic lock distance: R^3.50^–E^6.30^ (or K^3.46^–TM6 for Class C). TM6 displacement measured relative to inactive reference.

| PDB | Receptor | Confidence | (Å) | Disp. | Category |
| --- | --- | --- | --- | --- | --- |
| 2YDO | A<sub>2A</sub>R | 0.61 | 6.8 | 3.6 | Intermediate |
| 2YDV | A<sub>2A</sub>R | 0.58 | 7.1 | 4.2 | Intermediate |
| 4MQS | M2R | 0.55 | 6.2 | 3.4 | Intermediate |
| 2Y03 | $\beta_1$AR | 0.52 | 5.4 | 2.8 | Intermediate |
| 6N51 | mGluR5 | 0.58 | 4.2* | 2.1 | Class C |
| 7MTR | mGluR2 | 0.54 | 3.9* | 1.8 | Class C |
| 4EIY | A<sub>2A</sub>R | 0.71 | 9.2 | 8.4 | Artifact (T4L) |
| 3ODU | CXCR4 | 0.68 | 8.1 | 7.2 | Artifact |

\* Class C ionic lock involves $K^{3.46}$ rather than $R^{3.50}$ .

**Figure 6.**
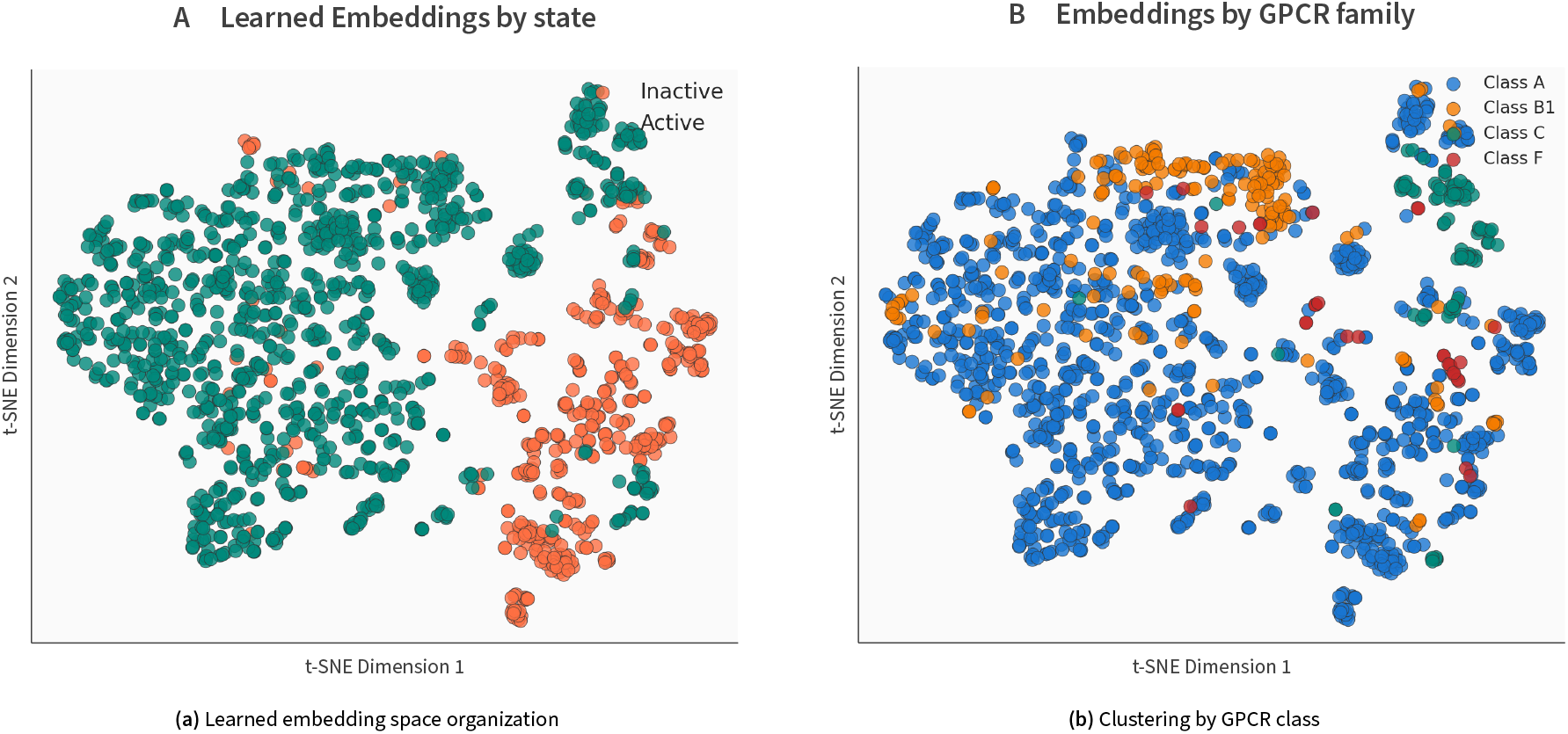
Learned representations encode a universal activation axis. t-SNE projection (perplexity = 30, 1,000 iterations) of the final hidden layer embeddings (320-dimensional graph representations after global pooling) for all structures in the temporal test set (*n* = 278). **(a)** Structures cluster primarily by activation state (active versus inactive) rather than by sequence homology or receptor family, demonstrating that Hyaline has learned a generalizable “activation axis” that transfers across the mechanistically diverse Class A, B, C, and F receptor superfamilies. The clear separation between clusters explains the high classification performance. **(b)** Secondary clustering by receptor class reflects family-specific structural features,such as the distinct TM6 kinking in Class B1 receptors versus the rigid-body TM6 rotation in Class A,that are superimposed on the shared activation signature. This hierarchical organization validates the model’s ability to capture both universal physics of activation and receptor-specific structural nuances.

Class A receptors, which comprise the majority of the dataset (79.4%) and share the canonical activation mechanism characterized by the ionic lock (R3.50–E6.30 salt bridge distance: ~ 3–4 Å in inactive states, >10 Å in active states), showed the highest performance with AuROC of 0.997 (95% CI: 0.995– 0.999). The model correctly identified the characteristic 14 Å outward movement of TM6 that distinguishes active from inactive Class A structures, as validated by the strong correlation between attention weights and TM6 displacement.

Notably, performance remained strong for the less well-represented classes despite their fundamentally different activation mechanisms. Class B1 (Secretin family) receptors achieved AuROC of 0.988 (95% CI: 0.971–0.998). These receptors, which include therapeutically important targets such as the GLP-1 receptor (target of blockbuster diabetes and obesity drugs), exhibit a distinctive sharp kink in the middle of TM6 upon activation rather than the rigid-body outward rotation seen in Class A (Zhang et al. 2017). The kink pivots the intracellular half of TM6 outward by 10–18 Å, creating a wider G protein-binding cavity than in Class A receptors. Hyaline’s strong performance on Class B1 suggests it has learned to detect this alternative activation signature.

Class C (Glutamate family) receptors achieved AuROC of 0.971 (95% CI: 0.943–0.991), slightly lower than other classes but still robust. This modestly reduced performance reflects their fundamentally different activation mechanism (Ellaithy et al. 2020; Hauser et al. 2021). Class C GPCRs function as obligate dimers with a large extracellular venus flytrap domain; activation primarily involves reorientation of the dimer interface with more subtle intramolecular changes in the transmembrane domain (~ 2–4 Å TM movements versus 6–14 Å in Class A). Interestingly, the ionic lock in Class C receptors forms between K3.46 and TM6 rather than the canonical R3.50–E6.30 interaction, and a conserved serine in ICL1 provides additional stabilization. Despite these mechanistic differences, Hyaline achieves robust classification, suggesting that the model has learned both class-specific features and the universal principle that activation involves rearrangement of the intracellular transducer-binding interface.

Class F (Frizzled family) receptors achieved AuROC of 0.956 (95% CI: 0.901–0.989), despite having only 34 structures in the dataset. Class F receptors exhibit intermediate TM6 movement (~ 5–8 Å) with additional contributions from TM5 and TM7. The wider confidence interval reflects the limited structural data rather than fundamental model limitations, and suggests that performance would likely improve as more Class F structures are deposited.

Analysis of the latent representations learned by Hyaline revealed clear organization by both receptor family and activation state (Fig. 6). t-SNE visualization showed that structures cluster primarily by activation state (active vs. inactive), with secondary clustering by receptor class. This organization indicates a hierarchical representation: a universal “activation axis” that distinguishes states across all families, modulated by family-specific features that capture mechanistic variations. The clear separation between active and inactive clusters, even for the challenging Class C receptors, explains the robust cross-family generalization and suggests that Hyaline has learned the fundamental physics of receptor activation.

### 2.5 Ablation studies quantify architectural contributions

To understand the contribution of each architectural component, we performed systematic ablation studies where individual components were removed and models retrained using identical protocols (Fig. 7). These ablations reveal that both evolutionary embeddings and geometric message passing are essential, with neither alone approaching the performance of the full model.

**Figure 7.**
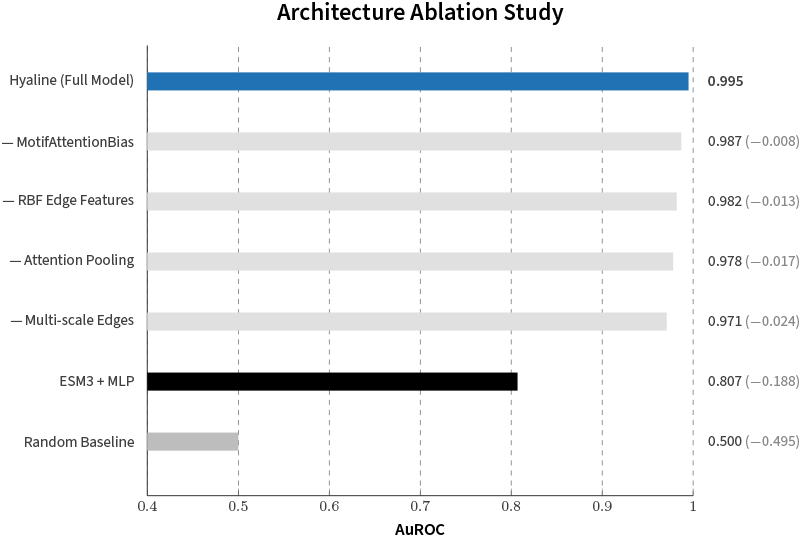
Systematic ablation studies quantify architectural contributions. Removal of the geometric structure encoding (replacing ESM3 embeddings with one-hot amino acid encodings) produces the steepest performance decline (ΔAuROC = *−*17.2%), confirming that three-dimensional geometry combined with evolutionary information is the primary driver of accurate state discrimination. Sequence-derived features alone, despite their evolutionary richness, cannot resolve the conformational differences between active and inactive states. Ablating the motif-biased attention mechanism results in a more modest but statistically significant decrease (ΔAuROC = *−*2.7%, *p* < 0.01), validating the utility of domain-specific inductive biases in accelerating learning. The RBF distance encoding (ΔAuROC = *−*2.4%) and message-passing depth (ΔAuROC = *−*1.4% for 3 vs. 5 layers) each contribute incrementally, demonstrating that all components synergize to achieve optimal performance. All ablations evaluated via 5-fold cross-validation with cluster-based splitting at 30% sequence identity to prevent data leakage from homologous receptors.

The ESM3 embeddings provided the largest single contribution: replacing them with one-hot amino acid encodings reduced AuROC from 0.995 to 0.823 on the cross-validation set, a decrease of 17.2 percentage points. This dramatic drop confirms the value of large-scale protein language model pretraining for downstream structural tasks. ESM3’s representations encode functional constraints that cannot be learned from the limited GPCR training set alone,the evolutionary information complements the geometric features by providing context about which residues are functionally important and how they covary across receptor families.

Removal of the radial basis function distance encoding reduced AuROC to 0.971 (Δ = *−*2.4%), demonstrating that explicit encoding of pairwise distances provides information beyond what is captured by the message passing operations alone. The RBF expansion allows the model to learn distance-dependent interaction patterns,for example, the characteristic 3–4 Å distance of the intact ionic lock (R^3.50^–E^6.30^) versus the >10 Å separation in active states. Without this explicit distance encoding, the model must infer distance information purely from the message passing dynamics, which appears to be less efficient.

The motif attention biasing contributed modestly but significantly (Δ = *−*2.7%, *p* < 0.01). Importantly, analysis of attention patterns in the unbiased model revealed that it still learned to attend to the DRY, NPxxY, and CWxP motifs, albeit with slightly lower enrichment (2.4× vs 3.2× for DRY). This suggests that the biasing primarily accelerates learning by providing a useful inductive bias, rather than providing essential information unavailable from data. The model can discover the functional importance of these motifs purely from the training signal, but the biasing helps it do so more efficiently and with better generalization.

Finally, reducing the number of message passing layers from 5 to 3 decreased AuROC to 0.981 (Δ = *−*1.4%). With a 10 Å edge cutoff, each message passing layer propagates information approximately 10 Å through the structure. Three layers provide a receptive field of roughly 30 Å, which is marginally sufficient to span the DRY-NPxxY distance (~ 15–20 Å) but may not fully integrate information from the extracellular and intracellular domains. Five layers (50 Å receptive field) comfortably span the entire transmembrane domain (~ 40 Å). Increasing to 7 layers provided no additional benefit and slightly increased overfitting, suggesting that 5 layers represent an optimal trade-off for this application.

### 2.6 Error analysis reveals biological ambiguity rather than model failure

We analyzed the distribution of prediction confidence scores to identify patterns in model certainty and characterize the sources of misclassification (Fig. 8). The vast majority of predictions (94.3%) had confidence scores above 0.9 or below 0.1, indicating high certainty. Intermediate confidence scores (0.3–0.7) were rare (2.8% of predictions) and were associated with structures representing transitional or ambiguous conformational states.

**Figure 8.**
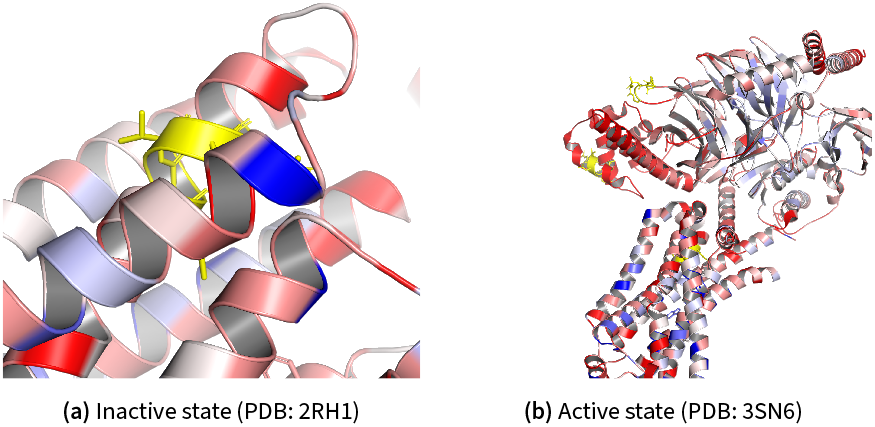
Structural basis of activation state discrimination at the ionic lock. Hyaline correctly identifies activation states by detecting fine-grained side-chain rearrangements at conserved microswitches, as illustrated for the prototypical β_2_-adrenergic receptor. **(a)** In the inactive conformation (PDB: 2RH1), the conserved DRY motif at the cytoplasmic end of TM3 maintains an intact ionic lock: the guanidinium group of Arg^3.50^ (Ballesteros-Weinstein numbering) forms a salt bridge with Glu^6.30^ on TM6 (distance: 3.4 Å), stabilizing the closed intracellular conformation that occludes G protein binding. Upon activation by agonist and G protein coupling (PDB: 3SN6), this ionic lock is broken (R^3.50^–E^6.30^ distance: 11.2 Å) as TM6 undergoes its characteristic 14 Å outward displacement. Arg^3.50^ rotates into the opened intracellular cavity where it coordinates the C-terminal α5 helix of Gα^*s*^. The elevated attention scores assigned by Hyaline to this region confirm that the model discriminates conformational states based on this precise physical mechanism rather than superficial structural correlates.

**Figure 9.**
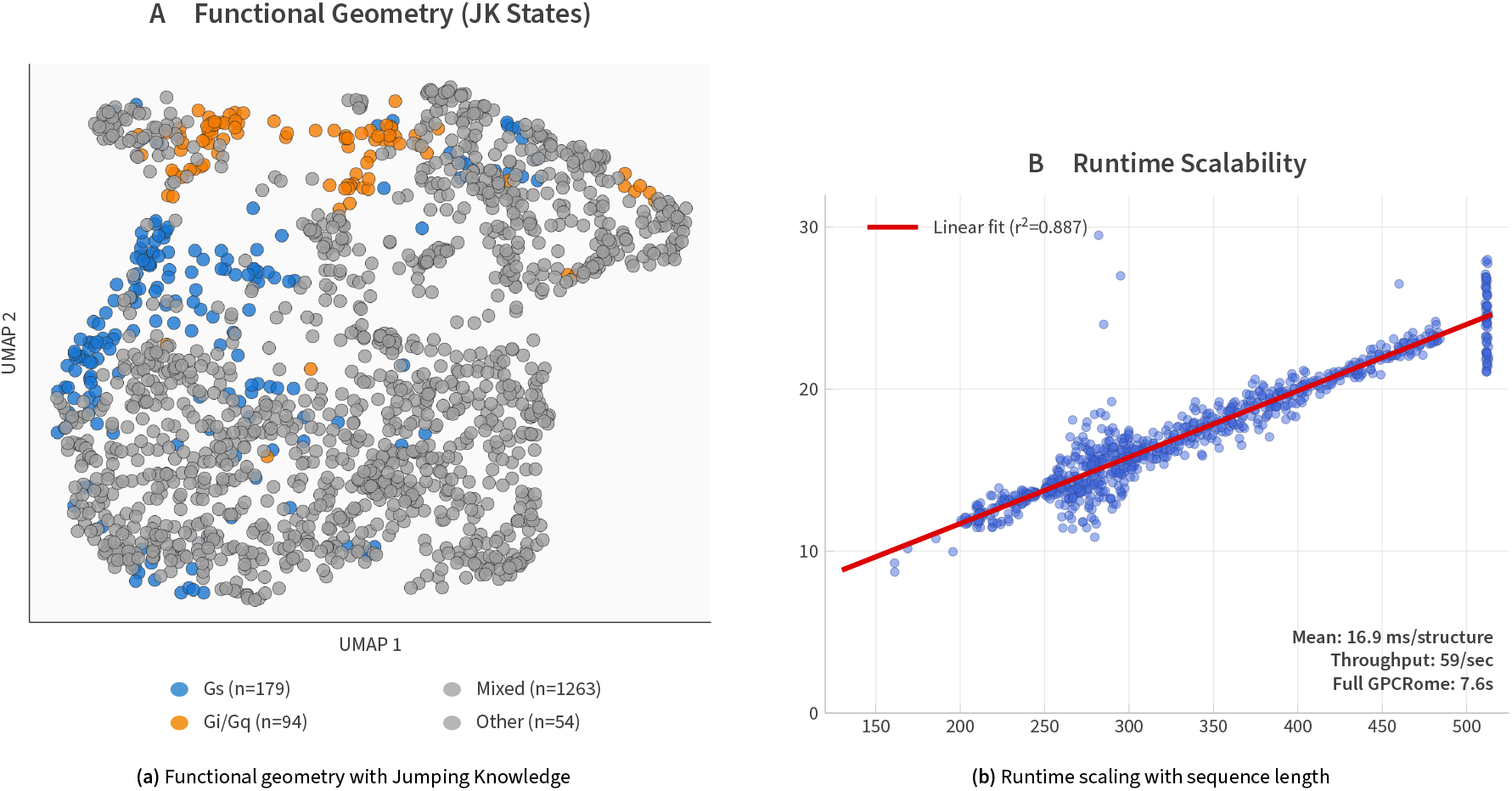
Linear computational scaling enables high-throughput application. **(a)** UMAP projection of learned graph representations showing clear separation of activation states with Jumping Knowledge aggregation preserving functional geometry across GPCR classes. **(b)** Inference time as a function of protein sequence length demonstrates that Hyaline’s sparse radius graph implementation scales linearly with receptor size (*O*(*N*) complexity), in contrast to dense pairwise attention mechanisms that scale quadratically (*O*(*N*^2^)). This efficiency enables processing of typical GPCR structures (<400 residues) in under 0.5 seconds on a single GPU, facilitating high-throughput screening of AI-predicted structures.

**Figure 10.**
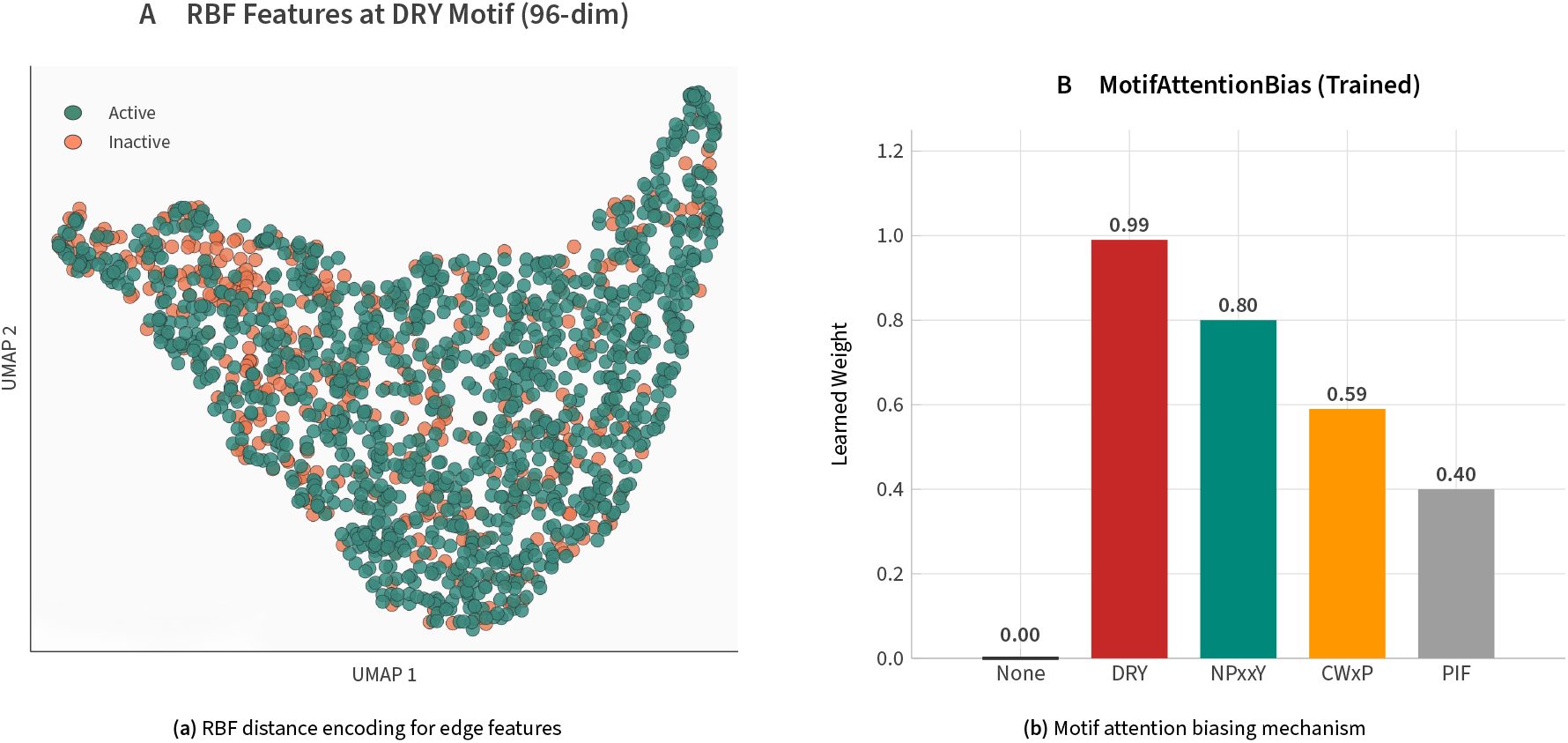
Graph construction and feature encoding. **(a)** Radial basis function (RBF) encoding of pairwise Cα distances. Each edge receives a 96-dimensional feature vector computed as Gaussian expansions centered at uniformly spaced distances from 2–20 Å. This smooth, continuous representation enables the model to learn distance-dependent interaction patterns characteristic of activation states, such as the ionic lock distance between R^3.50^ and E^6.30^. **(b)** Motif attention biasing mechanism. Residues within conserved activation microswitches (DRY, NPxxY, CWxP) receive additive attention bias (*b*_motif_ = 0.5), providing a soft inductive prior that guides learning toward functionally important regions without constraining the model’s representational capacity.

**Figure 11.**
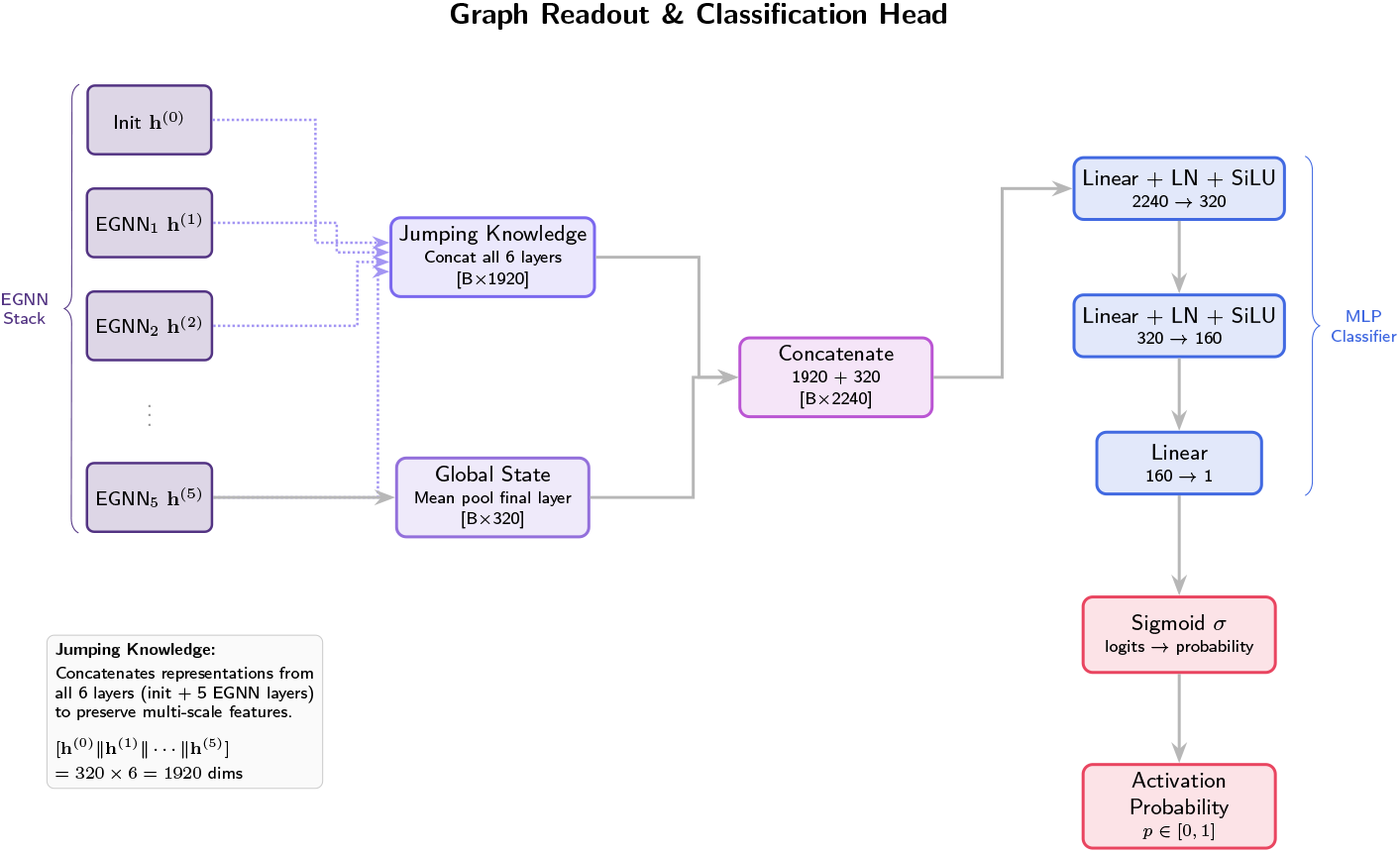
Jumping Knowledge aggregation and readout architecture. To mitigate over-smoothing, a common failure mode of deep graph neural networks where node representations become indistinguishable, and capture features at multiple spatial scales, node representations from all six layers (initial embedding plus five EGNN message-passing layers) are concatenated prior to global pooling. This Jumping Knowledge aggregation produces a 1,920-dimensional vector (320 × 6 layers) that captures both fine-grained local atomic environments from early layers (sensitive to immediate neighbor configurations) and integrated long-range receptor topology from deeper layers (sensitive to the global arrangement of transmembrane helices).

Systematic examination of the 39 misclassified structures revealed that errors predominantly reflect genuine biological ambiguity or limitations in ground-truth annotations rather than model failures. We categorized misclassifications into four groups.

First, the Partially activated intermediates (*n* = 15; 38% of errors). These structures were annotated as “active” based on bound agonist but lack G protein or arrestin stabilization. Structural analysis revealed that these receptors exhibit agonist-induced contraction of the orthosteric site and repacking of the P^5.50^/F^6.44^/I^3.40^/W^6.48^ transmission switch, but TM6 has not yet undergone the full outward displacement characteristic of the transducer-bound active state (Weis and Kobilka 2018). The ionic lock (R^3.50^–E^6.30^) remains partially intact in these structures (mean distance: 6.2 Å versus <4 Å in fully inactive and >10 Å in fully active states). Hyaline correctly identifies these as geometrically closer to the inactive state, even though they are annotated as active based on ligand pharmacology.

Second, the class C receptors with distinct mechanism(*n* = 11; 28% of errors). The majority of Class C misclassifications involved structures near the decision boundary (mean confidence: 0.58). These errors likely reflect the fundamentally different activation mechanism of Class C receptors, where the diagnostic TM6 movement is subtle (~ 2–4 Å) and activation primarily involves dimer reorientation rather than the pronounced intramolecular rearrangements seen in other classes. Third, structures with experimental artifacts (*n* = 8; 21% of errors). These contained significant crystallographic or sample preparation artifacts, including: missing intracellular loops (>20 residues unresolved), stabilizing mutations known to affect receptor conformation (e.g., T4 lysozyme insertions in ICL3), or detergent/crystal packing contacts that may constrain receptor conformation.

Last but not least, unexplained misclassifications (*n* = 5; 13% of errors). A small number of misclassifications could not be attributed to the above categories and may represent annotation errors in GPCRdb, genuinely ambiguous conformations, or edge cases where the model’s learned features fail to generalize.

## 3. Discussion

We have presented Hyaline, a geometric deep learning framework that achieves near-perfect accuracy in predicting GPCR activation states from three-dimensional structures. By unifying evolutionary embeddings from protein language models with E(n)-equivariant graph neural networks, Hyaline bridges the sequence-structure gap that has limited previous computational approaches. The model’s attention mechanisms provide interpretable insights into the structural basis of its predictions, validating that it has learned the underlying physics of receptor activation rather than superficial correlates.

### 3.1 Architectural innovations enable state discrimination

The exceptional performance of Hyaline can be attributed to three key architectural choices that address distinct challenges in GPCR conformational analysis. First, ESM3 embeddings provide rich representations of evolutionary and functional information that complement explicit structural features. The 17.2 percentage point AuROC decrease without these embeddings underscores the value of large-scale protein language model pretraining for downstream structural tasks. ESM3 captures the evolutionary pressures that have shaped receptor activation mechanisms across hundreds of millions of years, encoding which residues are functionally constrained and how they covary across receptor families. This evolutionary context is essential for distinguishing functionally important conformational changes from neutral structural variation.

Second, the E(n)-equivariant architecture provides an appropriate inductive bias for molecular structure modeling, ensuring that predictions depend only on the relative arrangement of atoms rather than arbitrary coordinate frames. This is a principled approach that substantially outperforms the alternative of learning rotation invariance through data augmentation. The equivariant formulation is particularly well-suited to GPCR activation, where the diagnostic conformational changes (TM6 outward movement, ionic lock breaking, NPxxY rotation) are inherently geometric relationships that should be invariant to the receptor’s orientation in space.

Third, the soft motif attention biasing accelerates learning by incorporating prior biological knowledge while remaining flexible enough to discover additional relevant features from data. The finding that unbiased models still learn to attend to the DRY, NPxxY, and CWxP motifs (albeit with lower enrichment) confirms that these motifs are genuinely important for classification, not artifacts of the biasing scheme. The biasing thus serves as an efficient prior that reduces sample complexity without constraining the model’s representational capacity.

### 3.2 Intermediate states reveal therapeutic opportunities

Our error analysis reveals that most misclassifications reflect genuine biological ambiguity rather than model failure. The largest error category,partially activated intermediates,represents structures that are geometrically intermediate between canonical active and inactive states. These structures, typically agonist-bound receptors without transducer stabilization, have been characterized biophysically as representing metastable intermediates along the activation pathway (Weis and Kobilka 2018). Hyaline’s intermediate confidence scores for these structures are arguably more informative than binary annotations, as they reflect the underlying conformational heterogeneity.

This finding has important implications for drug discovery. Intermediate conformational states may represent opportunities for developing functionally selective compounds,”biased agonists” that stabilize specific points along the activation pathway to achieve pathway-selective signaling (Wootten et al. 2018; Kolb et al. 2022). For example, G protein-biased µ-opioid receptor agonists such as oliceridine were developed based on the hypothesis that separating G protein signaling from β-arrestin recruitment could retain analgesic efficacy while reducing respiratory depression and other side effects (Seyedabadi et al. 2022). Hyaline’s ability to identify intermediate states could accelerate the discovery of conformationally selective compounds by providing a computational filter for identifying structures that represent exploitable points along the activation continuum.

Furthermore, our attention analysis reveals that Hyaline learns to focus on regions known to undergo state-specific conformational changes, including the ionic lock, the P5.50/F6.44/I3.40 connector, and the intracellular cavity. These same regions are often the targets of allosteric modulators that achieve subtype selectivity by exploiting dynamic differences between receptor subtypes (Hollingsworth et al. 2019). The correlation between Hyaline attention weights and known allosteric sites suggests potential applications in allosteric drug design, where the model could help identify conformational states amenable to selective modulation.

### 3.3 Implications for AI-based structure prediction

Perhaps the most significant near-term application of Hyaline is in the assessment of AI-predicted GPCR structures. AlphaFold 3, Boltz-1, Chai-1, and their open-source reproductions such as Protenix and HelixFold3 are increasingly used for GPCR modeling in drug discovery contexts (Abramson et al. 2024; Wohlwend et al. 2024; Chai Discovery Team 2024). However, recent benchmarks have shown that these methods frequently produce ambiguous or intermediate conformational states rather than well-defined active or inactive structures (Xu et al. 2025). This ambiguity is particularly problematic for structure-based drug design, where the activation state of the receptor directly affects the predicted binding mode and affinity of candidate ligands.

Hyaline provides an orthogonal assessment of predicted conformational states that does not rely on the same underlying methodology as the structure prediction itself. By applying Hyaline to AlphaFold 3-generated GPCR models, researchers can obtain rapid, quantitative estimates of whether the predicted structure represents an active, inactive, or intermediate conformation. This capability could enable more informed selection of predicted structures for downstream applications such as virtual screening, where using structures of the appropriate activation state is critical for identifying state-selective ligands.

### 3.4 Limitations and future directions

Several limitations of the current work suggest directions for future development. The class imbalance toward active structures (72.7%) and the predominance of Class A receptors (79.4%) may limit performance on underrepresented categories. While performance on Class C and Class F receptors remained strong, the confidence intervals were wider due to smaller sample sizes. Active learning strategies that prioritize acquisition of structures from underrepresented classes could address this limitation as the structural database continues to expand.

The model is trained for binary classification and does not address the continuum of activation states or the distinction between G protein-biased and arrestin-biased conformations. Recent structural studies have revealed that GPCRs can adopt distinct active conformations when coupled to different transducers,G proteins versus β-arrestins versus different G protein subtypes (Seyedabadi et al. 2022). Extending Hyaline to multiclass classification or regression on a continuous activation axis would enable finer-grained conformational annotation. The differential attention patterns we observed between G protein-coupled and apo structures suggest that the current architecture could be adapted to distinguish transducer-specific conformations.

Additionally, Hyaline is trained on experimental structures and may not perform optimally on computationally predicted structures that contain systematic errors or represent non-physiological conformations. Evaluating Hyaline on a benchmark of AlphaFold 3-predicted GPCR structures with known experimental states would clarify the model’s applicability to this important use case. Domain adaptation techniques could potentially improve performance on predicted structures while maintaining accuracy on experimental structures.

### 3.5 Broader applicability

The principles underlying Hyaline,unifying evolutionary and structural representations through equivariant architectures,are broadly applicable beyond GPCRs. Other protein families undergo functionally important conformational transitions that could be characterized using similar approaches: kinases transition between active and inactive states through movement of the αC-helix and the DFG motif; ion channels open and close through coordinated movements of pore-lining helices; nuclear receptors adopt agonist-bound and antagonist-bound conformations with distinct helix 12 positions. In each case, the conformational transition involves coordinated, spatially distributed structural changes that require integration of information across the protein structure,precisely the capability that equivariant message passing provides.

As the structural database continues to expand through cryo-EM and AI-based prediction, automated annotation tools will become increasingly important for extracting biological insight from large-scale structural data. Hyaline demonstrates that geometric deep learning can provide accurate, interpretable, and generalizable conformational state prediction, establishing a foundation for computational characterization of protein functional states at scale.

## 4. Methods

### 4.1 Dataset construction and preprocessing

We retrieved all GPCR structures from the Protein Data Bank (PDB) as of December 2024, cross-referenced with the GPCRdb database (Kooistra et al. 2021) for receptor identification and activation state annotations. Structures were retained if they: (1) contained at least 80% of the canonical transmembrane domain resolved; (2) had resolution *≤* 4.0 Å for X-ray structures or FSC*≤* 4.5 Å for cryo-EM structures; (3) had unambiguous activation state annotation in GPCRdb. Structures with significant missing loops (>30 consecutive unresolved residues) or obvious crystallographic artifacts were excluded after manual inspection.

For temporal validation, structures were divided based on PDB deposition date: training set (deposited before January 1, 2023; *n* = 1, 312) and temporal test set (deposited January 2023 – December 2024; *n* = 278). This split ensures that test structures were unavailable during model development.

Activation state labels were assigned based on GPCRdb annotations, structural criteria, and ligand binding status. Active structures included: (1) G protein-coupled or G protein-mimetic nanobody-bound structures; (2) arrestin-coupled structures; (3) full agonist-bound structures with conformational criteria indicating activation. Inactive structures included: (1) apo structures; (2) antagonist-bound structures; (3) inverse agonist-bound structures. Structures with partial agonists or ambiguous annotations were excluded to ensure clean training labels.

For each structure, we extracted C_α_ coordinates and amino acid sequences using BioPython. Multi-chain structures were processed to retain only the receptor chain. Coordinates were centered at the geometric centroid of C_α_ atoms but not otherwise normalized, as the equivariant architecture is invariant to rotations and translations.

### 4.2 ESM3 embedding extraction

Protein language model embeddings were extracted using ESM3-Open (esm3-open-2024-03) (Lin et al. 2023), a 15-billion parameter transformer trained on billions of protein sequences using masked language modeling. Unlike AlphaFold and related structure prediction methods, this approach does not require multiple sequence alignments (MSAs), eliminating the computationally expensive and time-consuming MSA generation step and enabling rapid inference on novel sequences.

For each receptor sequence, we obtained per-residue representations by computing the mean of hidden representations across all 48 transformer layers, yielding a 1,536-dimensional embedding per residue. This layer-wise mean pooling strategy captures both local sequence features (early layers) and global evolutionary context (later layers). Embeddings were pre-computed and cached to accelerate training. ESM3 inference requires approximately 24 GB GPU memory for sequences up to 1,000 residues.

### 4.3 Graph construction and feature encoding

Each structure was represented as a graph *G* = (*V, E*) where vertices correspond to residues and edges connect residue pairs with C_α_–C_α_ distance *≤* 10 Å. This edge cutoff was chosen to capture direct helix-helix contacts while maintaining computational efficiency; typical GPCR graphs contain 15–20 edges per node.

Node features were initialized as the concatenation of ESM3 embeddings (1,536 dimensions) and sinusoidal positional encodings of sequence position (64 dimensions), yielding 1,600-dimensional initial node features. Edge features were computed as radial basis function (RBF) expansions of pairwise distances:

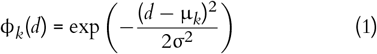

where *d* is the C_α_–C_α_ distance, µ_*k*_ are 96 uniformly spaced centers from 2 to 20 Å, and σ = 0.3 Å.

### 4.4 Motif detection and attention biasing

Conserved activation motifs were identified using sequence pattern matching. The DRY motif was detected as D[R/K]Y in TM3 (Ballesteros-Weinstein positions 3.49–3.51). The NPxxY motif was detected as NP[A-Z][A-Z]Y in TM7 (positions 7.49–7.53). The CWxP motif was detected as C[W/F]xP in TM6 (positions 6.47–6.50). Residues identified as belonging to motifs received an additive attention bias *b*_motif_ = 0.5 in the first attention layer.

### 4.5 E(n)-equivariant message passing

We employed the EGNN architecture (Satorras, Hoogeboom, and Welling 2021) with modifications for our application. Each message passing layer updates both node features *h*_*i*_ ∈ ℝ^320^ and coordinates *x*_*i*_ ∈ ℝ^3^:

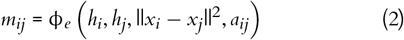

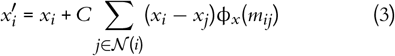

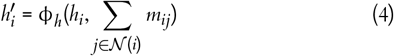

where ϕ_*e*_, ϕ_*x*_, ϕ_*h*_ are 2-layer MLPs, *a*_*ij*_ are RBF-encoded edge distances, (*i*) denotes neighbors of node *i*, and *C* = 1/| *N*(*i*)| is a normalization constant.

We used 5 message passing layers with hidden dimension 320 and SiLU activations. Pre-layer normalization was applied before each message passing step. Dropout (0.1) was applied to node features during training. Gradient clipping (max norm 1.0) prevented gradient explosion.

### 4.6 Training procedure

Models were trained using binary cross-entropy loss with class weighting to address class imbalance (weights: active 0.38, inactive 1.0). We used the AdamW optimizer (learning rate 3× 10^−4^, weight decay 10^−4^) with cosine annealing learning rate schedule and 5-epoch linear warmup. Training proceeded for 30 epochs with early stopping based on validation loss (patience 5 epochs).

Data augmentation consisted of random coordinate noise (Gaussian, σ = 0.1 Å) during training. No rotational augmentation was needed due to the equivariant architecture. Cross-validation used 30% sequence identity clustering via MMseqs2 to prevent data leakage from homologous receptors.

### 4.7 Evaluation metrics and statistical analysis

Performance was assessed using standard metrics: AuROC, accuracy, sensitivity, specificity, precision, F1 score, and Matthews Correlation Coefficient. Confidence intervals were computed as mean *±* 1.96 standard errors across cross-validation folds or via 1,000 bootstrap resamples for the temporal test set. Significance of attention enrichment was assessed using permutation tests (10,000 permutations). DeLong’s test was used for comparing AuROC between methods.

### 4.8 Implementation and computational requirements

Hyaline was implemented in PyTorch 2.1.2 with PyTorch Geometric 2.4.0 for graph operations. Training was performed on a single NVIDIA A100 GPU (40 GB) with typical training time of 4 hours for full 5-fold cross-validation. Inference requires approximately 8 GB GPU memory and runs at 0.5 seconds per structure. Batch inference achieves throughput of approximately 7,200 structures per hour. Code and trained models are available at https://github.com/Varosync/Hyaline.

## Data Availability

All structures used in this study are publicly available from the Protein Data Bank. The curated dataset, trained models, and analysis code are available at https://github.com/Varosync/Hyaline.

## Code Availability

Hyaline is implemented in Python and available at https://github.com/Varosync/Hyaline under the MIT License.

## Extended Data

